# Bio-inspired artificial printed bioelectronic cardio-3D-cellular constructs

**DOI:** 10.1101/2022.01.26.477866

**Authors:** Paola Sanjuan-Alberte, Charlie Whitehead, Joshua N. Jones, João C. Silva, Nathan Carter, Simon Kellaway, Richard J.M. Hague, Joaquim M.S. Cabral, Frederico C. Ferreira, Lisa J. White, Frankie J. Rawson

## Abstract

Bioelectronics is a growing field where novel smart materials are required to interface biology with electronic components. Conductive hydrogels have recently emerged as a promising material for biosensing/actuating applications as they can provide a wet, nanostructured and electrically conductive environment, minimising the mismatch between biological and electronic systems. In this work, we propose a strategy to develop conductive bioinks compatible with the freeform reversible embedding of suspended hydrogels (FRESH) extrusion bioprinting method. These bioinks are based on decellularized extracellular matrix (dECM), extracted from three different tissues (small intestine submucosa, liver and bone) and were characterised. 3D structures were manufactured containing human pluripotent stem cell-derived cardiomyocytes (hPSC-CMs), exhibiting cell viabilities >80%. Multi-walled carbon nanotubes (MWCNTs) were selected as an additional component of the bioinks. The addition of the MWCNTs enhanced the conductive features of the hydrogels and the morphology of the dECM fibres. Electrical stimulation (ES) through alternating currents was applied to hPSC-CMs encapsulated in 3D structures manufactured with the previous material and our results indicated two main findings: (1) in the absence of external ES, the conductive properties of the materials can improve the contractile behaviour of the hPSC-CMs and (2) this effect is significantly enhanced under the application of external ES. Genetic markers analysed showed a trend towards a more mature state of the cells evaluated by the TNNI3/TNNI1 ratio, with upregulated SERCA2 and RYR2 calcium handling proteins when compared to controls and downregulation of calcium channels involved in the generation of pacemaking currents (CACNA1H). These results demonstrate the potential of our strategy to manufacture conductive hydrogels in complex geometries for bioactuating purposes. However, further development of the 3D bioprinting techniques is required to achieve higher control over the nano- and microarchitectures of the structures to improve their biomimicry.

## 1. Introduction

Cell-material interactions have traditionally been one of the main focuses of biomaterials and tissue engineering research. In the last decade, there has been an increased demand for smart and stimuli-responsive materials to provide additional control over material’s properties and cell fate [1]. Enhanced functionalities are particularly important to improve the biomimicry of electroconductive tissues. For instance, it has been now widely accepted that conductive environments promote neural proliferation and differentiation [2, 3]. Additionally, the development of bioelectronic systems and devices relies on the interface between biological and electroconductive systems.

Despite the increased popularity of synthetic conductive substrates for their ability to influence cell behaviour, conductive natural biomaterials represent a better alternative due to their tissue-like characteristics and mechanical properties [4, 5]. There is also a mismatch in the conductive mechanism between electrically conductive synthetic substrates and electroconductive tissues that needs further addressing and investigation [6]. This mismatch can be minimised using conductive hydrogels, as these provide an ion-rich and wet physiological environment in a three-dimensional (3D) nanostructured and conductive network [7]. The electrical conductivity of hydrogels can be increased by the incorporation of conductive micro- and nanofillers within the hydrogel matrix [8]. It has been hypothesised that the incorporation of the conductive materials bridges the insulating pore walls of the hydrogels, propagating the electrical signals and stimulating cell constructs evenly and uniformly [9]. The most commonly conductive nanofillers used include metallic nanoparticles [10], conductive polymers [11] and carbon-based nanomaterials [12]. In bioelectronics, conductive hydrogels have been explored for the development of wearable electronics [13], implantable devices [14] and sensing/actuating applications [15].

There is a wide variety of natural biomaterials used for the development of conductive hydrogels. Decellularized extracellular matrix (dECM) materials have shown promise because functional and structural components of native ECM can be retained [16, 17], maintaining the biochemical cues that naturally interact with cells in a specific microenvironment [18]. Furthermore, dECM can be used in the composition of bioinks that subsequently allow the additive manufacturing of 3D structures [19]. dECM-based hollow tubes and bifurcating structures resembling anatomical features such as blood vessels and airways have been 3D printed using the freeform embedding of suspended hydrogels (FRESH) extrusion method [20]. The variety of tissues from which dECM can be extracted determines the versatility and functionality of the bioprinted structures, where intrinsic cellular morphologies and functions can be reconstituted [21]. There have been two recent reports on making dECM conductive with addition of carbon-based nanomaterials and subsequently merging them with cardiomyocytes [22, 23]. However, neither included detailed analysis of effect of the material in the electrical genotype of the cells. Such analysis is important as we have previously suggested that this is one of the most important functions to modulate when aiming at in vitro generation of mature cardiomyocytes [24].

In this work, a conductive bioink for the 3D bioprinting of structures has been developed, combining the electroconductive features of multi-walled carbon nanotubes (MWCNTs) with the biochemical and structural cues of dECM. Such inks have never been explored in 3D bioprinting. Initially, a general strategy to formulate inks and bioinks for FRESH extrusion bioprinting based on dECM extracted from several tissues was established. Once this was achieved, electroconductive hydrogels were formulated and characterised. Finally, and to explore the bioelectronic applications of this material, electrical stimulation to 3D printed structures containing cardiac cells was performed, evaluating the potential of the materials to regulate cardiac cells’ fate.

## 2. Materials and methods

### Tissue decellularization and characterisation

All the tissues used in this work were harvested from 12-to 24-month-old animals from an EU-certified butcher and/or abattoir.

For the preparation of the small intestine submucosa extracellular matrix (sisECM), porcine small intestines were abundantly washed with water and intestinal contents were discarded. Using forceps and a razor blade, the intestine was cut and opened and the serosa, mucosa and muscularis layers were removed, leaving behind the submucosa layers [25]. the resultant tissue was rinsed with water and cut into smaller pieces, which were then washed in distilled water (dH_2_O) under agitation for 2 h. Tissue depyrogenation was carried out in peracetic acid (PAA) (Acros Organics) 0.1% in a 24:1 dH_2_O/ethanol solution for 2 h under agitation and washed with phosphate buffer saline (PBS) and dH_2_O.

Decellularized liver extracellular matrix (lECM) was prepared from a previously frozen porcine liver. Frozen tissue was sliced into 3-4 cm cuts and washed in dH_2_O to remove any excess blood in a mechanical shaker. This solution was then replaced with a 0.02% Trypsin (Sigma-Aldrich)/0.05% EDTA (Fischer Scientific) solution in PBS, incubated at 37°C for 1 h. A 3% Triton-X-100 (Sigma-Aldrich) in dH_2_O was then used for 1 h at room temperature. Finally, tissues were washed with 4% deoxycholic acid (Sigma-Aldrich) (w/v) in dH_2_O [26]. In between solution exchanges, tissues were thoroughly washed with dH_2_O and pressed between pieces of mesh to aid in cell lysis [26]. PAA depyrogenation was carried out following the aforementioned procedure.

Decellularized bone extracellular matrix (bECM) was prepared following previous protocols [27, 28] from cancellous bone segments. Briefly, bone segments were washed with 0.1% w/v gentamicin in PBS. Following this, segments were immersed in liquid nitrogen and ground in a bladed grinder. Bone granules were demineralised in 0.5 N HCl (25 mL per gram of bone) for 24 h at room temperature under stirring and washed with dH_2_O. Lipids were extracted using a 1:1 mixture of chloroform/methanol (30 mL per gram of bone) for 1 h at room temperature and washed with methanol and dH_2_O. Enzymatic decellularization was performed using a 0.05% trypsin (Sigma-Aldrich)/0.02% EDTA (Fisher Scientific) solution in PBS at 37°C for 24 hours under constant agitation. The decellularized bone was then washed in an equal volume of PBS at 37°C for 24hrs under constant agitation, and then rinsed in dH_2_O.

In all cases, the decellularized materials were lyophilised, milled and stored at – 20°C for further use.

Quantification of DNA and sulphated glycosaminoglycans was performed as described previously [29]. Briefly, the determination of decellularization efficiency was based on quantification of dry weight double-stranded DNA (dsDNA). Concentrations of dsDNA were measured using a Quant-iT™ Picogreen® assay kit (Thermo Fischer #P7589) following the instructions provided by the supplier. Sulphated GAGs were determined using a 1,9 dimethyl-methylene blue (DMMB) assay. A papain buffer consisting on 100 mM of Na_2_HPO_4_, 10 mM of Ethylenediamine Tetraacetic Acid (EDTA), 10 mM of glycine and 125 μg mL^-1^ papain was used to digest dECM at a concentration of 10 mg mL^-1^. The previous materials were provided by Sigma-Aldrich. A blank papain buffer was used as a negative control and diluent for the assay. A total of 50 μL of samples were mixed with 200 μl of DMMB solution (0.03 M sodium formate, 0.046 M DMMB, ethanol [0.5% v/v] and formic acid [0.2% v/v] in a 96-well plate. The standard curve ranged from 0 to 100 μg mL^-1^ chondroitin-4-sulphate. Absorbance was measured at 525 nm and 595 nm. Samples are analysed by taking the difference of the readings between absorbance at 525nm and 595nm (OD_525_–OD_595_).

### dECM ink formulation and printing

Lyophilised dECM material was enzymatically digested in a solution of 1 mg mL^-1^ porcine pepsin (Sigma-Aldrich #P7012) in 0.01 N HCl under stirring for 48 h at room temperature to obtain dECM digest with a final concentration of 10 mg mL^-1^. These solutions were stored at -20°C when not immediately used. To induce the gelation of the dECM digests (pH = 2-3), a neutralisation buffer consisting of 0.1 M NaOH (Sigma-Aldrich) and PBS was added to the dECM digest following previous protocols [28] and kept at 4°C to prevent the spontaneous gelation before the 3D printing process was carried out, with a final concentration of 8 mg mL^-1^. Prior to printing, this material was loaded into a 3cc QuantX™ syringe barrel with a 27G straight cannula blunt end dispensing tip (Fisnar). This pre-hydrogel solution was kept at 4°C during the preparation and printing process to prevent its gelation before the printing process was carried out.

In samples containing -COOH functionalised multi-walled carbon nanotubes (MWCNTs) (>95%, OD: 10-20 nm, US Nano) these were dispersed at a concentration of 5-10 mg mL^-1^ in the neutralisation buffer and mixed with the dECM digests as described previously to reach a final concentration of 1-2 mg mL^-1^ in the dECM pre-hydrogel solution. A schematic of this can be seen in **Figure S1**.

For the 3D bio-/printing, a custom microextrusion-based 3D printing system was used as described previously [30]. Briefly, the system consisted of a three-axis dispensing robot (Fisnar F5200N) and a pneumatic dispensing unit (Ultimus V, Nordson EFD) interfaced with a personal computer. Printing speeds for the dECM bioinks ranged 0.8-2 mm s^-1^ and the printing pressures used ranged 0.1-0.5 psi. Coordinates of the dispensing robot were programmed using G-code commands. The gelatine support bath was prepared using a FRESH Kit and LifeSupport™ for FRESH (Allevi #FF-0002) in a Well-plate.

After printing the different structures, printed samples were kept at room temperature for 20 minutes to induce the gelation of the dECM and then incubated at 37°C for 30 minutes to melt the gelatine support bath. Once the gelatine melted, it was carefully aspirated and structures were kept in PBS or culture media.

### Gelation kinetics and stability over time of printed dECM structures

A BioTek ELx800 plate reader interfaced with a personal computer incorporated with the Gen5 2.06 data analysis software was used to measure the absorbance of the dECM hydrogels over time. The absorbance was measured at 450 nm (at room temperature) and measurements were repeatedly undertaken for 60 min at 2 min intervals on the neutralised digested dECM. Values were normalised to allow further comparison between the different tissues. Data were normalised using **Eq.1** and the turbidity profile was fitted to a sigmoid curve to calculate the t_1/2_, the time needed to reach 50% of the maximum turbidity absorbance value, and slope. The slope determined the speed of gelation. Statistical significance was calculated using a paired t-test.

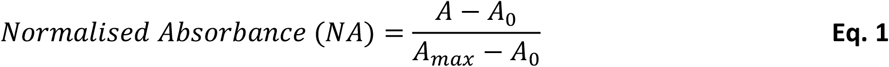

Where is A is absorbance.

For stability studies, 6 mm side squares and 6 mm diameter rings with three layers each were printed using the aborementioned conditions and incubated in PBS. Images of the structures were obtained on days 0, 3, 5, 7, 10, 15, 20, 30 and 60 post-printing.

### hPSC-CMs differentiation and bioink formulation

A REBL-PAT hPSC line was derived from a skin punch biopsy from a male subject. Procedures of isolation, culture, differentiation and dissociation are described elsewhere [31]. Dissociation of cells took place 6–8 weeks after the differentiation process. Cell culture media consisted in basal RPMI medium (Life Technologies #11875093) supplemented with B27 (Life Technologies #17504044, Carlsbad, CA, USA), Y-27632 ROCK inhibitor (20 μM; Tocris #1254) and 10% foetal bovine serum (FBS, Sigma-Aldrich). The medium was changed after 24 h to RPMI/B27 medium. Early hPSC-CMs were incorporated into the bioinks by adding the cell suspension (1/5 of the final volume of the bioink) at an approximate concentration of 0.5-1 million cells mL^−1^ to the neutralised dECM pre-hydrogels. Bioprinting was carried out using the aforementioned parameters.

### hPSC-CMs purity assessment

hPSC-CMs were fixed in 2% paraformaldehyde (VWR) for 30 min at room temperature and permeabilised with 0.1% Triton-X 100 (Sigma-Aldrich) for 8 min at room temperature. Non-specific binding was blocked with 4% FBS (Sigma-Aldrich) in PBS for 1 h at room temperature. Cells were then immunostained with a monoclonal primary antibody against TNNI3 produced in mice (Sigma-Aldrich # WH0007137M4) at a concentration of 1:1000 in PBS overnight at 4°C. A solution of 0.05% of Tween-20 was used to wash the samples and a solution of goat anti-Mouse secondary antibody IgG was added (1:1000, Abcam # ab6785) and incubated for 2 h at room temperature. Samples were washed with the previous Tween-20 solution and exposed to Hoechst 33258 (5 μg mL^-1^, Sigma-Aldrich). Fluorescence images were taken on a fluorescence microscope (Leica DMI 3000B equipped with Nikon-AcT1 software). The purity of differentiated cells was analysed by calculating the ratio of hPSC-CMs (cells stained both with Hoechst and TNNI3) vs the total population of cells by fluorescence microscopy. Two different batches of cells were analysed in triplicate and two images were quantified per sample. Statistical significance was calculated using a t-test (N=3, n=2; ±SD)

### hPSC-CMs viability assays

Viability studies were performed on bioprinted 10 × 10 mm squared meshes using the different bioinks. On day 7, structures were carefully washed in PBS and incubated for 30 min in 1 μM acetoxymethyl (AM) calcein solution (Sigma Aldrich #C1359) in PBS to stain viable cells (green) and dead cells (red) were stained with 5 μM ethidium homodimer I (Sigma Aldrich #E1903) in PBS. Fluorescence images were taken in the aforementioned fluorescence microscope. Images were processed using the ImageJ open-source software (https://imagej.nih.gov/ij/). To assess the effects of shear stress on hPSC-CMs viability during the printing process, different pressures (1, 2, 5 and 10 psi) and needle diameters (200 μm, 400 μm and 600 μm OD) were studied on cell suspensions in culture media and compared to control conditions (hPSC-CMs cultured on a polystyrene well-plate). Experiments were performed in triplicate.

### Rheological characterisation

A Physical MCR 301 (Anton Paar, UK) instrument was used for the rheological characterisation of the pre-gels. For this time sweep analysis, 200 μL of the samples were added to a pre-cooled to 4°C Peltier plate. A Peltier hood was used to keep the humidity conditions stable during the measurements. The geometry used was a 25 mm diameter parallel plate at a working distance of 0.4 mm. After samples were in place, the temperature of the Peltier plate was increased to 37°C. Angular frequency and amplitude were kept constant at 1 rad s^-1^ and 1%, respectively. Readings were taken once every 15 s for 20 min to obtain storage (G’) and loss (G’’) modulus. The gelation point was then calculated, corresponding to the moment where G’ = G’’

Amplitude sweep analyses were performed immediately after the previous test. The angular frequency remained constant at 1 rad s^-1^ and the oscillatory strain ramped from 1-200%. The temperature was kept constant at 37°C.

For each test, four samples were analysed per material (n=4). Statistical significance was calculated using one-way ANOVA.

### Semi-quantification of printability

Semi-quantification of the printability of bECM, bECM+MWCNTs 1mg mL^-1^ and bECM+MWCNTs 2 mg mL^-1^ was calculated following previously described protocols [32]. Briefly, 10 × 10 mm square meshes were 3D printed with the different materials and the perimeter of the square gaps of the mesh was calculated using ImageJ. The printability was then calculated using the following formula:

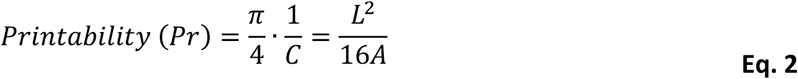

Where *C* corresponds to circularity, *L* to the length of the perimeter and *A* to the area of the perimeter.

### Electrochemical characterisation of conductive hydrogels

A Keithley 2450 source meter (Tektronix) connected to a four-probe station was used for the conductivity measurements of dried samples. 10 mm lines of bECM, bECM+MWCNTs 1 mg mL^-1^ and bECM+MWCNTS 2 mg mL^-1^ were 3D printed and dried at room temperature. A fixed current of 10 μA was applied through the outer probes and the resulting potential was measured at the inner probes. Readings were taken once every 1 s for 30 minutes. From the stable region of the graphs, the sheet resistance was then calculated (n=3, ±SD).

Electrochemical impedance spectroscopy (EIS) was performed on a PGSTAT potentiostat including the FRA32M module (Metrohm Autolab, The Netherlands) and interfaced with a personal computer including the NovaLab software. The electrochemical cell consisted of two sputter-coated gold electrodes (Georg-Albert PVD, Germany) separated by a 2 mm gap in a two-electrode configuration at room temperature. Hydrogels were casted using moulds with 2 mm height and 10 mm diameter and sandwiched between the gold electrodes. Frequencies applied ranged between 10-100000 Hz and the amplitude was 0.01 V_RMS_.

### Swelling degree

For the assessment of the swelling degree, 6 mm diameter rings of the different materials were 3D printed and immersed in liquid nitrogen followed by lyophilisation. The initial weight of the dried materials was recorded (W_d_) and then samples were immersed in PBS at 37°C. At specific time points (1, 5, 10, 15, 20, 30, 45, 60, 90 and 120 min), the water excess was carefully removed using Kimtech paper and the weight of the samples was recorded (W_w_) (n=3). The swelling percentage was calculated using the formula below:

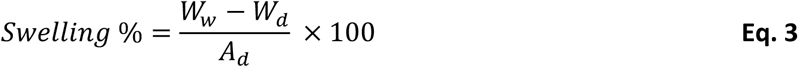

### Scanning electron microscopy imaging

6 mm diameter rings were printed and mounted on filter paper to facilitate the manipulation of the samples. Before SEM imaging, samples were dried using a Samdri-780a (Tousimis, US) critical point drier. 100% ethanol was used before CO_2_ addition. After drying, samples were mounted on an aluminium pin stub with carbon tape prior to coating with a 5 nm layer of iridium on a Leica EM ACE 600. SEM images were obtained on a field-emission scanning electron microscope (FEG-SEMl JOEL 7100F) in secondary electron mode with an acceleration voltage of 5 kV and a working distance of 10 mm. Fibre diameters were calculated by measuring the diameter of 100 fibres per sample and histograms were produced using GraphPad Prism v8.4.3.

### Electrical stimulation and measurement of cellular contractility

Bioprinted structures of bECM and bECM+MWCNTs 1 mg mL^-1^ containing hPSC-CMs at a concentration of 1 million cells mL^-1^ were electrically stimulated for five days during 2 h. Electrical stimulation was provided using an AFG1022 arbitrary function generator (Tektronix). Stimulation parameters consisted of a square wave with an amplitude of 4 V (2 V to -2 V) at a frequency of 1 Hz. Quantification of the contraction rate was calculated from videos of the hPSC-CMs on the different samples with and without electrical stimulation using the MYOCYTER v1.3 macro for Image J. This macro enables the scaling of the time-dependent changes of pixel intensity in subsequent frames of the recorded cardiomyocytes, enabling the depiction of cellular contractility as positive amplitudes on an arbitrary scale. Data extraction was performed according to the developer’s instructions [33] (N=3, n=2, ±SD). Statistical significance was calculated using a t-test.

### Gene Expression analysis

hPSC-CMs were dissociated from bioprinted structures and total RNA was isolated using the RNeasy Mini kit (Qiagen) following the manufacturer’s guidelines. RNA concentrations were calculated using a NanoVue Plus spectrophotometer (GE Healthcare). RNA was reverse transcribed using the High Capacity cDNA Reverse Transcription Kit (Thermo Fischer Scientific) and a T100™ thermal cycler (Bio-Rad). The expression of selected genes was determined by RT-PCR using the iQ Universal SYBR Green Supermix (Bio-Rad) and a StepOnePlus real-time PCR equipment (Applied Biosystems). Gene expression was normalised to the housekeeping gene GAPDH and results were analysed using the 2^-ΔΔCt^ method (N=3, n=3, ±SD). Statistical significance was calculated using one-way ANOVA. The specific primer sets (StabVida) used are listed in **Table S1**.

## 3. Results and Discussion

### Tissue decellularization and bioink formulation

Tissue decellularization was successfully achieved from three different organs: porcine small intestine submucosa (sisECM), porcine liver (lECM) and bovine cancellous bone (bECM). Our choice was based on the fact that these organs are readily available and the dECM extraction protocols have been validated previously [27, 34, 35], providing versatile and robust protocols for bioink formulation and 3D bioprinting with these materials. These materials have also been previously reported in the literature for applications in cardiac tissue repair [36, 37]. In the case of sisECM, cells were removed by mechanical delamination. The native tissue can be seen in **Figure 1a** and the results of the decellularization in **Figure 1b**. In the case of lECM, the dECM extraction process involved enzymatic and chemical removal of the cells with detergents and images of native and lECM can be seen in **Figure 1c** and **Figure 1d**, respectively. For bECM, the process included demineralisation and delipidation prior to enzymatic decellularisation, with images of native and bECM shown in **Figure 1e-f**.

**Figure 1.**
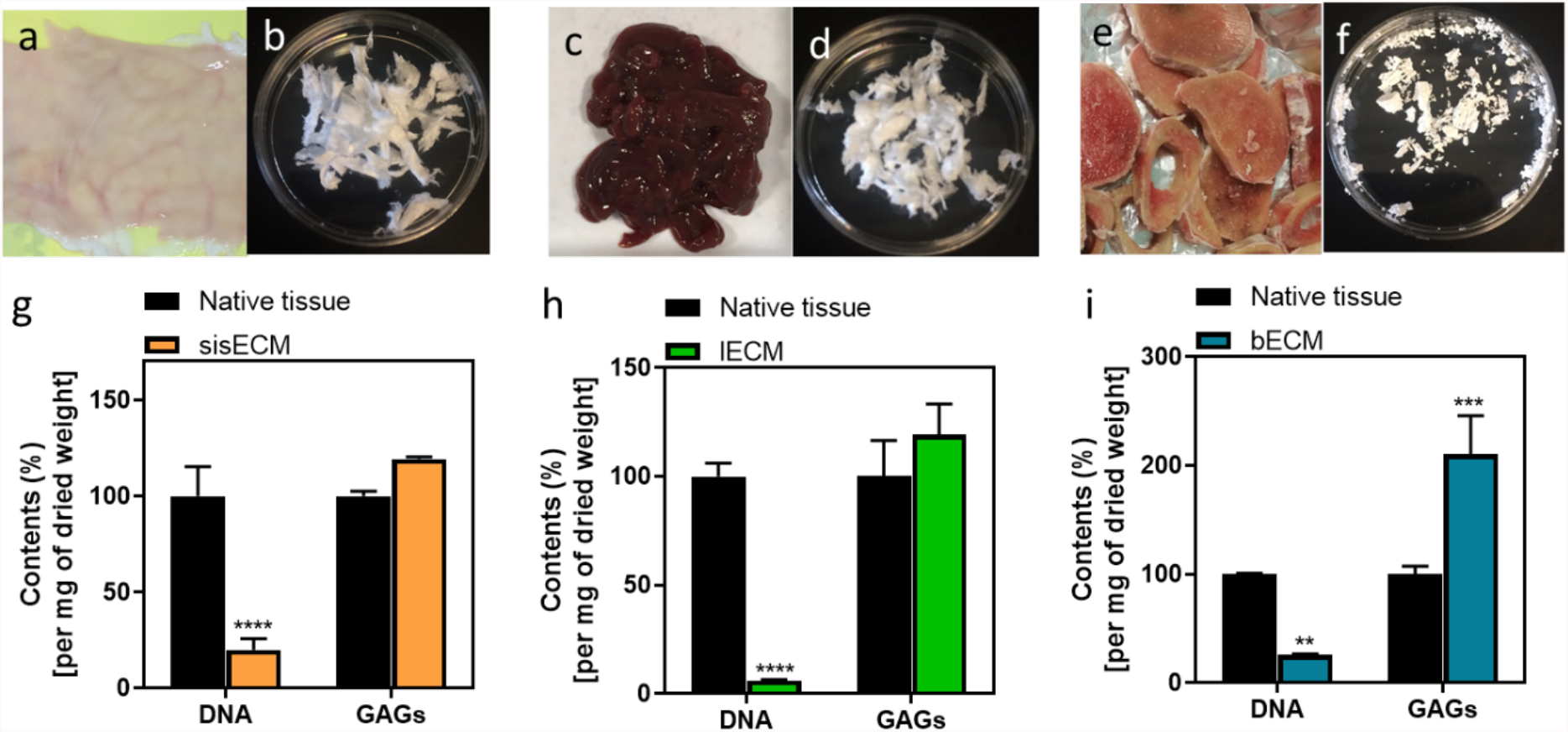
Three organs were decellularized for extracellular matrix (ECM) extraction. Porcine small intestine submucosa (sis) **(a)** before and **(b)** after decellularization (sisECM). Porcine liver **(c)** before and **(d)** after decellularization (lECM). Bovine bone **(e)** before and **(f)** after decellularization (bECM). Percentage of DNA and glycosaminoglycans (GAGs) present in native and decellularized ECM in **(g)** sis, **(h)** liver and **(i)** bone. Quantification was performed per mg of dried tissue. Composition of native tissue were assumed as 100% (n=3).

The main purpose of the decellularization process is to remove the native cells from these tissues while preserving the ECM structure and composition, which is not a trivial task [38]. This is because residual cellular material can induce cytotoxic effects when ECM biomaterials are implanted *in vivo*. The amount of residual cellular material present in the decellularized tissues can be quantified by calculating the amount of DNA present in the dECM as remnant DNA can be directly correlated with residual cells within the dECM [39]. Although the main goal is to remove cells effectively, the ECM structure and components such as collagen and glycosaminoglycans (GAGs) need to be preserved during the decellularization process. The quantification of structural molecules is therefore crucial to evaluate the quality of the decellularized products.

In the case of DNA quantification, the DNA content of the dECM is significantly reduced after the decellularization process as expected, corresponding to 19.84% for sisECM, 6.05% for lECM and 25.98% for bECM (**Figure 1g-1i**, respectively). The lower DNA percentage was obtained in lECM as a combination of enzymatic and chemical methods to remove cells are usually more effective. From these results we concluded that the tissues have been effectively decellularized and the structural molecules have been preserved.

In all tissues, the GAG content was relatively high when compared to native tissues, indicating that the decellularization process did not cause any structural damages to the extracted dECM (**Figure 1g-1i**). The GAG quantification was normalised per mg of dried tissue and the contents of the native tissues were assumed as 100%. Since the native tissues still preserve the cellular material, the weight of the ECM is more diluted, causing the GAG content of the dECM to be >100%, similar to that described previously [21].

### FRESH extrusion bio-/printing

The FRESH extrusion printing method was used in the manufacturing of complex dECM structures. This method consists of the printing of materials inside a gelatine slurry and is commonly used for the bio-/printing of hydrogels as it allows the deposition and cross-linking of soft biomaterials while avoiding their collapse and deformation during the printing process [40] (**Figure 2a**). Once structures were printed, the gelation process of the printed material was induced *in situ* by increasing the temperature to room temperature, inducing the spontaneous collagen fibrillogenesis process. The gelatine support bath was later removed by increasing the temperature to 37°C, which will lead to the melting of gelatine microparticles and consequently to the release of the printed structures. An example of the complexity of the structures manufactured using the bECM ink can be seen in **Figure 2b**. The advantage of the FRESH technique over conventional extrusion bioprinting methods [41] of dECM-based bioinks is that no photo-crosslinking with UV is required.

**Figure 2.**
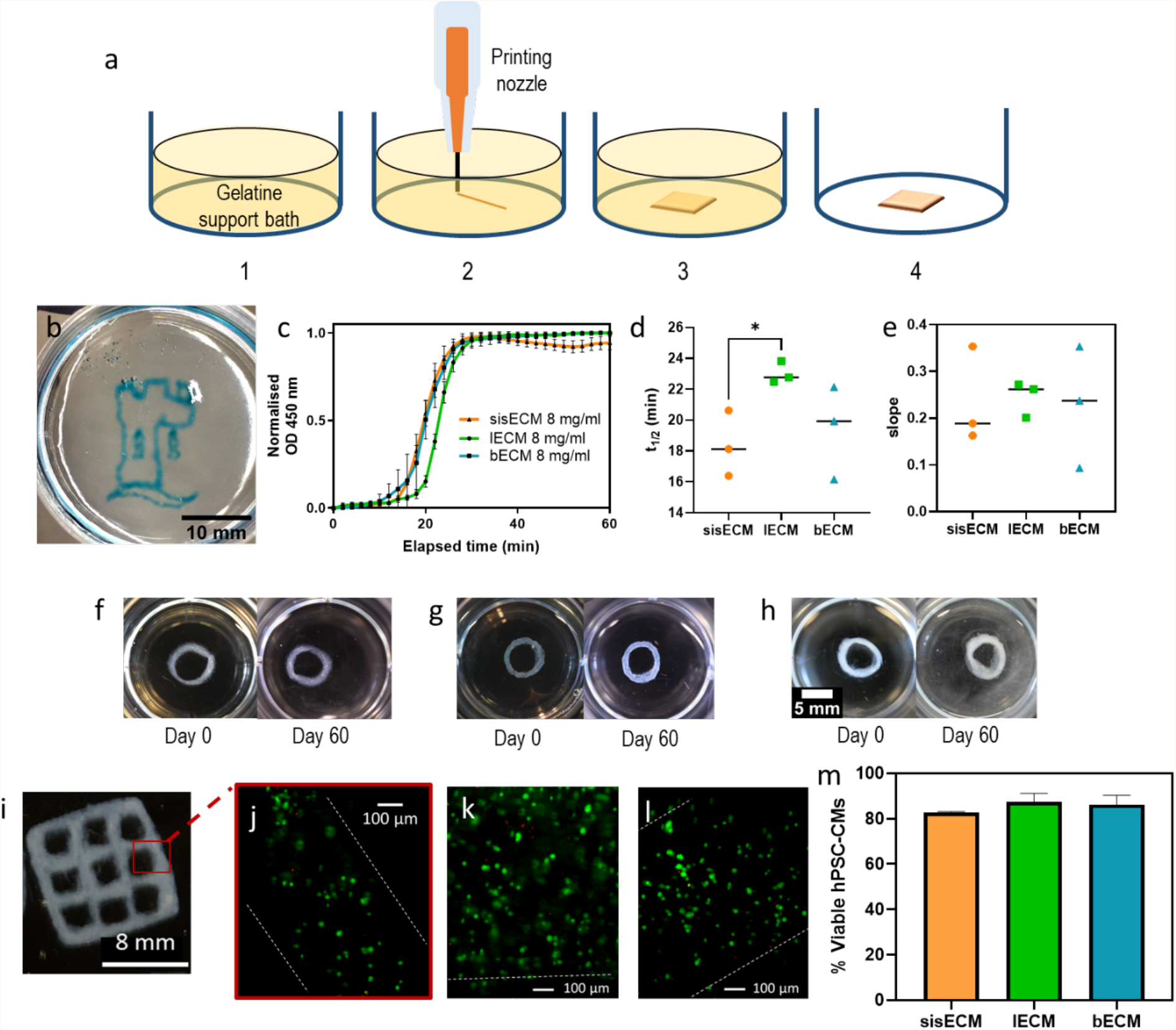
**(a)** Schematic representation of the process of FRESH extrusion printing of ECM hydrogels. 1. A thermo-reversible support bath formed by gelatine microparticles is used as a substrate. 2. Extrusion printing of cold decellularized ECM (dECM) takes place inside the gelatine bath. 3. *In situ* gelation of printed dECM structures at room temperature. 4. Structure is released when the temperature is increased to 37°C. **(b)** Printed bECM structure following the FRESH extrusion method. Scale bar 10 mm. **(c)** Normalised turbidimetric gelation kinetics of sisECM, lECM and bECM at 450 nm (n=3). Determination of the kinetics parameteres **(d)** t_1/2_ and **(e)** slope. Five-layered printed 6 mm diameter rings of **(f)** sisECM, **(g)** lECM and **(h)** bECM and their appearance on day 0 and day 60 after printing. **(i)** Example of a 10 × 10 mm bECM 3D bioprinted scaffolds and inset of **(j)** fluorescence image of bioprinted hPSC-CMs in bECM after live/dead staining. Representative fluorescence microscopy images of bioprinted hPSC-CMs in **(k)** sisECM and **(l)** lECM after live/dead staining. **(m)** Percentage of viable cells after bioprinting using the different bioinks (n=3). The dotted white line indicates the edge of the structures. Images were taken on day 7 after bioprinting.

The intrinsic properties of the gels vary from tissue to tissue, therefore, the gelation kinetics of the different dECMs was evaluated to determine the time required for the structures to fully gelate after printing, allowing subsequent gelatine support bath removal. A turbidimetric evaluation was used on the different tissues, as changes in the turbidity of the solutions provide a rapid and reproducible means of monitoring the collagen fibrillogenesis [42]. As it can be seen in **Figure 2c**, the three distinct phases of fibrillogenesis can be observed: a lag phase, an exponential growth phase and a plateau phase. For the different dECM types, the turbidity profile was similar in all cases. The previous graphs were fitted into sigmoidal curves (**Figure S2**) to determine the kinetics parameters of t_1/2_, corresponding to the time needed to reach 50% of the maximum absorbance values, and the slope of the curves, corresponding to the rate of fibrillogenesis. Values of t_1/2_ of sisECM, lECM and bECM were 18.1, 22.8 and 19.9 min respectively, with a statistically significant difference between sisECM and lECM. For the slope, the data dispersion was bigger and no significant differences were observed. From this data, the faster gelation kinetics corresponded to sisECM, followed by bECM and lECM and we concluded that we could safely remove the gelatine support bath 30-40 min after the printing of the structures.

Printed dECM structures tend to be quite fragile and prone to damage due to their poor mechanical properties [43]. To assess the stability of the printed structures, 6 mm rings and squares were incubated in PBS for 60 days. As it can be seen in **Figure 2f-2h**, for all the three tissues, structures remained stable during the 60 days. Additional images can be found in **Figure S3**.

For the bioprinting experiments and to explore the bioelectronic applications of our materials, human induced pluripotent stem cell-derived cardiomyocytes (hPSC-CMs) were selected due to their electrogenic nature. These cells were differentiated from hPSCs following previous protocols [31] resulting in purities >95% of hPSC-CMs for the different batches (**Figure S4**). In order to optimise the bioprinting parameters and assess the shear stress effects on cell viability, suspended hPSC-CMs in culture media were extruded using different needles (200, 400 and 600 μm) and printing pressures (1, 2, 5 and 10 psi) (**Figure S5**). There were no observed major differences between the different conditions, with cell viabilities between 75-90% in all cases in contrast to the 90% viability observed at the controls (**Figure S6**). Some reduction in cell viability can be expected since hPSC-CMs are subjected to additional stress during the bioprinting process. Values >75% of viability are considered acceptable in bioprinting and are similar to other hPSC-CMs bioprinting studies [44, 45].

Once it was established that the bioprinting process does not affect in great measure the hPSC-CMs viability, hPSC-CMs were incorporated into the dECM bioinks. 10 mm^2^ meshes were bioprinted using the different dECM materials (**Figure 2i**). Calcein-AM and ethidium homodimer staining was performed 7 days after bioprinting of the structures and the results showed that high viability was maintained on the different bioinks (**Figure 2j-l**). It is important to note, that although some cells started to elongate, overall the spheroidal structure of the hPSC-CMs was maintained after bioprinting. This could be due to the lack of mechanical support offered by the dECM or cell-cell interactions, as hPSC-CMs are encapsulated in a 3D structure, as achieving elongated cells is currently one of the main challenges in bioprinting of cardiac tissues[46]. The percentage of viable hPSC-CMs was similar for the three types of dECM with the highest viability observed in lECM (87.3%) followed by bECM (86.2%) and sisECM (82.6%) (**Figure 2m**).

### Electroconductive dECM-based hydrogels

Multifunctional features were introduced in the dECM hydrogels by incorporating MWCNTs to the hydrogel formulations to increase their conductivity and explore the potential of this material for applications in bioelectronics. We have also previously observed that composites containing MWCNTs can affect the phenotype of hPSC-CMs [24], and thus, MWCNTs were selected as the conductive nano-filling material. To the best of our knowledge, composites of dECM and MWCNTs have never been previously used as bioinks in the bioprinting of tissues and their ability to modulate cell state has not been investigated.

To discard any cytotoxic effects associated to the incorporation of the MWCNTs, sisECM, lECM and bECM hydrogels containing 1 mg ml^-1^ MWCNTs (sisECM-MWNCTs, lECM-MWCNTs and bECM-MWCNTs, respectively) and a suspension of hPSC-CMs were casted using 5 mm moulds. Live/dead staining confirmed that most of the cells in the structures remained viable and that the incorporation of MWCNTs to the hydrogels did not induce noticeable cytotoxic effects in the hPSC-CMs (**Figure S7**).

Rheological characterisation of the different inks was performed in order to assess wether the effect of MWCNTs in the gelation and viscoelastic properties of the dECM materials could affect the printability of the inks. Initially, gelation kinetics were evaluated by increasing the temperature to 37°C during a time-sweep rheometric test in order to trigger the collagen fibrillogenesis process. As expected, both the storage and loss moduli of all samples increased, with rapid onset of gelation upon ramping the temperature to 37°C, indicating that the materials were transitioning to the gel state (**Figure 3a-c**). From these, it can be seen that bECM describes a more obvious sigmoidal curve than sisECM and lECM. The gelation point of plain sisECM, lECM and bECM was similar, with values of 1.63, 1.59 and 1.89 min respectively (**Figure 3d**). In the case of sisECM, the addition of the MWCNTs at concentrations of 1 mg mL^-1^ and 2 mg mL^-1^, did not significantly affect the gelation point of the inks. However, in lECM and bECM, the addition of the MWCNTs caused a decrease in the gelation time. Similar observations were also made in studies using carboxylic and hydroxyl-functionalised MWCNTs (MWCNTs-COOH and MWCNTs-OH) in glycol/chitosan gels [47] and MWCNTs-COOH in polysaccharide-based hydrogels [48]. One hypothesis to explain this observation could be that the carboxylic groups of the MWCNTs are contributing to the generation of additional bonds in the hydrogel.

**Figure 3.**
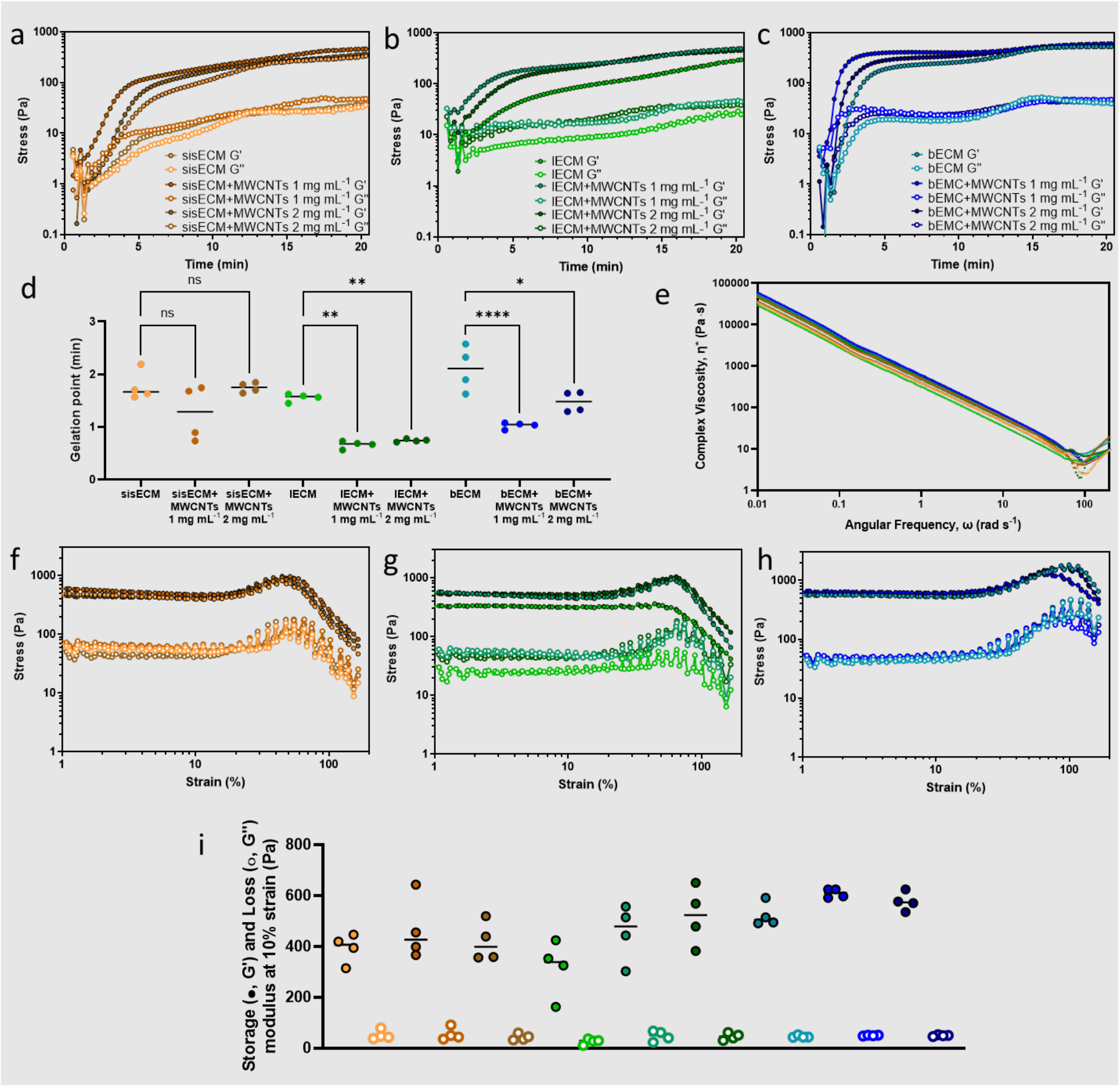
Rheological behaviour of the different dECM inks. Gelation kinetics showing the storage (G’) and loss moduli (G’’) over time of **(a)** sisECM, **(b)** lECM and **(c)** bECM with and without MWCNTs at 1 mg mL^-1^ and 2 mg mL^-1^. **(d)** Gelation point and **(e)** complex viscosity of the different materials. Strain-sweeps of **(f)** sisECM, **(g)** lECM and **(h)** bECM pre-gels with MWCNTs at 1 mg mL^-1^ and 2 mg mL^-1^ concentrations. **(i)** Values of storage (G’) and loss (G’’) modulus at 10% strain. Four samples were analysed for each hydrogel composition (n=4) from the same batch.

A frequency-sweep test was performed on all materials. Shear-thinning flow behaviour enables inks to be extrudable and reduce the shear forces exerted in the printing nozzles, and thus, a frequency-sweep test was performed in the materials. In all cases, the complex viscosity decreased linearly (**Figure 3e**), indicating a shear-thinning behaviour. At high values of frequency (> 50%) some irregularities can be seen in the graph, indicating some damage to the materials. Additionally, a strain-sweep test was also performed to evaluate the linear-viscoelastic (LVE) limit on the different materials once they transitioned to the gel state. In sisECM, a linear strain-stress behaviour up to 21% strain was observed (**Figure 3f**), where the addition of the MWCNTs did not seem to have a significant effect. In lECM the linear stress-strain region reached 44.5% strain (**Figure 3g**), and in the presence of MWCNTs, this value decreased to 25%. In bECM, linearity was observed up to 28% strain, with no noticeable differences amongst samples containing MWCNTs (**Figure 3h**). All samples exhibited decreasing G’ and G’’ values after approximately 50% strain, leading to catastrophic failures.

G’ and G’’ values from the different materials were compared at 10% strain, corresponding to the LVE region. G’ values of sisECM corresponded to 419 Pa (**Figure 3i**), and the addition of MWCNTs to the hydrogels did not seem to induce any changes to the material behaviour. In the case of lECM, G’ increases with the concentration of MWCNTs present in the gel, with values of 352 Pa, 443 Pa and 478 Pa for lECM, lECM+MWCNTs 1 mg mL^-1^ and lECM+MWCNTs 2 mg mL^-1^, respectively (**Figure 3i**). A similar trend was also observed in bECM, where G’ values were 491 Pa, 591 Pa and 576 Pa in bECM, bECM+MWCNTs 1 mg mL^-1^ and bECM+MWCNTs 2 mg mL^-1^ (**Figure 3i**). The magnitude of G’’ was similar in all samples and no noticeable differences were seen.

Despite all the materials exhibiting a similar rheological behaviour and presenting minor differences, it was not possible to process the sisECM and lECM inks containing MWCNTs in the bioprinter. We observed with these inks that continuous clogging of the needle tip was being produced, limiting considerably the printability of structures. For this reason, in subsequent bio-/printing experiments, bECM inks were selected. The printability (Pr) of these inks was evaluated on printed meshes with 10 × 10 mm. The semi-quantification of the Pr, was calculated based on previous works from Eq.2 [49], where the acceptable range of Pr was established at 0.9-1.1. In our case, the Pr of the bECM, bECM+MWCNTs 1 mg mL^-1^ and bECM+MWCNTs 2 mg mL^-1^ was < 1, but the values were within the acceptable printability region, and no significant differences were observed between them (**Figure S8**). The printed structures remained stable for up to 30 days (**Figure S9**).

Once it was determined that all the selected inks possessed acceptable printability values, we then proceeded to determine the resistivity and impedance values of the hydrogels using a 4-probe method and electrochemical impedance spectroscopy (EIS), respectively. For the determination of the resistivity values on the dry materials, 10 mm and 20 mm lines were printed and dried (**Figure 4a**). The results of the surface resistivity calculation indicate that the addition of the MWCNTs to the structures contributed to decrease the surface resistivity values, from 154 MΩ/sq in bECM to 106.1 MΩ/sq in bECM+MWCNTs 1 mg mL^-1^ and to 106.9 MΩ/sq in bECM+MWCNTs 2 mg mL^-1^ (**Figure 4b**). All resistivity values were below the controls (glass surfaces) (171 MΩ/sq).

**Figure 4.**
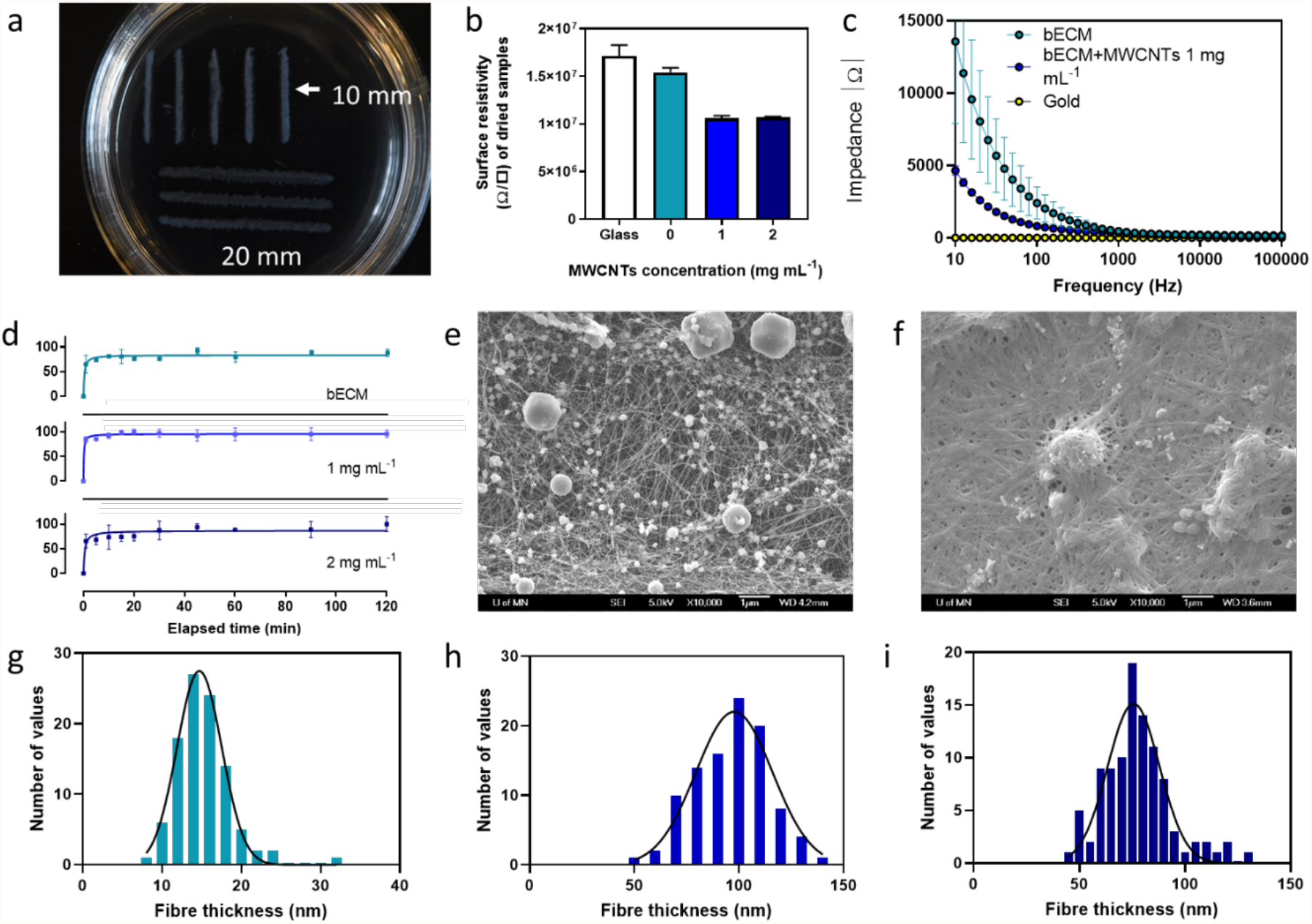
**(a)** 10 mm and 20 mm length printed structures used in the determination of surface resistivity. **(b)** Surface resistivity of dried printed samples (n=3). **(c)** Electrochemical impedance spectroscopy of bECM and bECM+MWCNTs 1 mg mL^-1^ compared to bare gold (n=3). **(d)** Swelling degree determination of printed bECM at increasing concentrations of MWCNTs (n=3). Scanning electron microscopy (SEM) images of **(e)** bECM and **(f)** bECM+MWCNTs 1 mg mL^-1^. Histogram analysis and Gaussian distribution of fibre thickness taken from SEM images of **(g)** bECM, **(h)** bECM+MWCNTs 1 mg mL^-1^ and **(i)** bECM+MWCNTs 2 mg mL^-1^ (n=100).

These results indicated the contribution of MWCNTs to decrease the resistivity of the materials. However, hydrogels are complex environments where both ionic and electronic currents and a synergy of these determine the overall conductivity of the materials. To investigate this, EIS measurements were conducted on hydrated samples, comparing bECM with bECM+MWCNTs 1 mg mL^-1^ and a fully conductive material (gold). On the samples containing MWCNTs, a decrease of an order of magnitude in values of impedance can be observed (**Figure 4c**), similarly to previous observations in gelatine methacrylate hydrogels containing MWCNTs [4, 50]. For instance, at 100 Hz, the values of impedance corresponded to 2408.7 in bECM, 823.1 in bECM+MWCNTs 1 mg mL^-1^ and 4.8 in gold, confirming that the addition of MWCNTs also contributes to increasing the conductivity of materials in wet conditions and the potential of this material in bioelectronics.

Further characterisation was performed on the bECM, bECM+MWCNTs 1 mg mL^-1^ and bECM+MWCNTs 2 mg mL^-1^ printed constructs to investigate any other contributions of the MWCNTs to the hydrogel structure. The swelling degree indicated a rapid hydration of the bECM lyophilised hydrogels, reaching a plateau phase after 20-30 min incubation (**Figure 4d**). In samples containing MWCNTs the recovery of the hydrogel structure followed the same trend.

Printed samples were also dehydrated using a critical point drying method to preserve their ultrastructure and imaged by SEM. These images showed significant differences in the morphology and thickness of the bECM fibres with and without MWCNTs (**Figure 4e-f**). Histogram analysis of the fibre thickness of the different samples was performed, where the average thickness of bECM fibres corresponded to 15.16 nm, in contrast to the 96.17 nm and 77.46 nm measured in bECM+MWCNTs 1 mg mL^-1^ and bECM+MWCNTs 2 mg mL^-1^, respectively. Interestingly, MWCNTs were not observed at these concentrations in contrast to images at lower MWCNTs concentrations of 0.2 mg mL^-1^, where bECM fibres and MWCNTs are easily distinguishable (**Figure S10**). Also, the typical collagen structure with a marked d-period was only observed at the higher MWCNTs concentrations. This suggests that an interaction between the bECM fibres and MWCNTs might be occurring at higher concentrations, leading to higher fibre diameters and reinforcing the strength of hydrogels as seen **Figure 3i**. It can also be noted from the SEM images that some salts are also present in the hydrogels. Higher magnification images can be seen in **Figure S11**.

### Effects of electrical stimulation (ES) on hPSC-CMs

The next part of the work was aimed at investigating the combination of biochemical cues, electrical conductivity, mechanical properties and ES on hPSC-CMs contractility and maturity. Previous studies demonstrated that hydrogels containing MWCNTs led to improved neuronal differentiation and morphology, and the differences were magnified under ES [51].

Bioprinted structures containing hPSC-CMs were subjected to a regime of 2 h per day for 5 days of ES (a square wave, 2 V to -2 V, 1 Hz) and the contractile behaviour of the cells was analysed. In bECM and no ES, the spontaneous contractions of the cardiomyocytes were either not observed or very sporadic (**Figure 5a, Figure S12, Video S1**). The contractility of bioprinted cells improved slightly in bECM+MWCNTs 1 mg mL^-1^ hydrogels, however, contractions still exhibit an erratic pattern typical of arrhythmic cardiac tissues (**Figure 5b, Video S2**). Under the application of ES, the contractions of the hPSC-CMs were more defined and rhythmic (**Figure 5c-d, Video S3**) and the contraction rate was significantly higher than in structures that were not subjected to ES (**Figure 5e, Video S4**), which was particularly enhanced in the presence of MWCNTs reaching physiological values. These results demonstrate two findings: (1) the material’s conductive properties alone can support improvements in hPSC-CMs contractile behaviour and; (2) such improvement on the contractile behaviour can be significantly enhanced when ES is applied. We tentatively hypothesise that this could be due to a combined effect of electrochemical and structural cues provided by the MWCNTs which mean the structures act as bipolar electrodes localising electric field effects[52].

**Figure 5.**
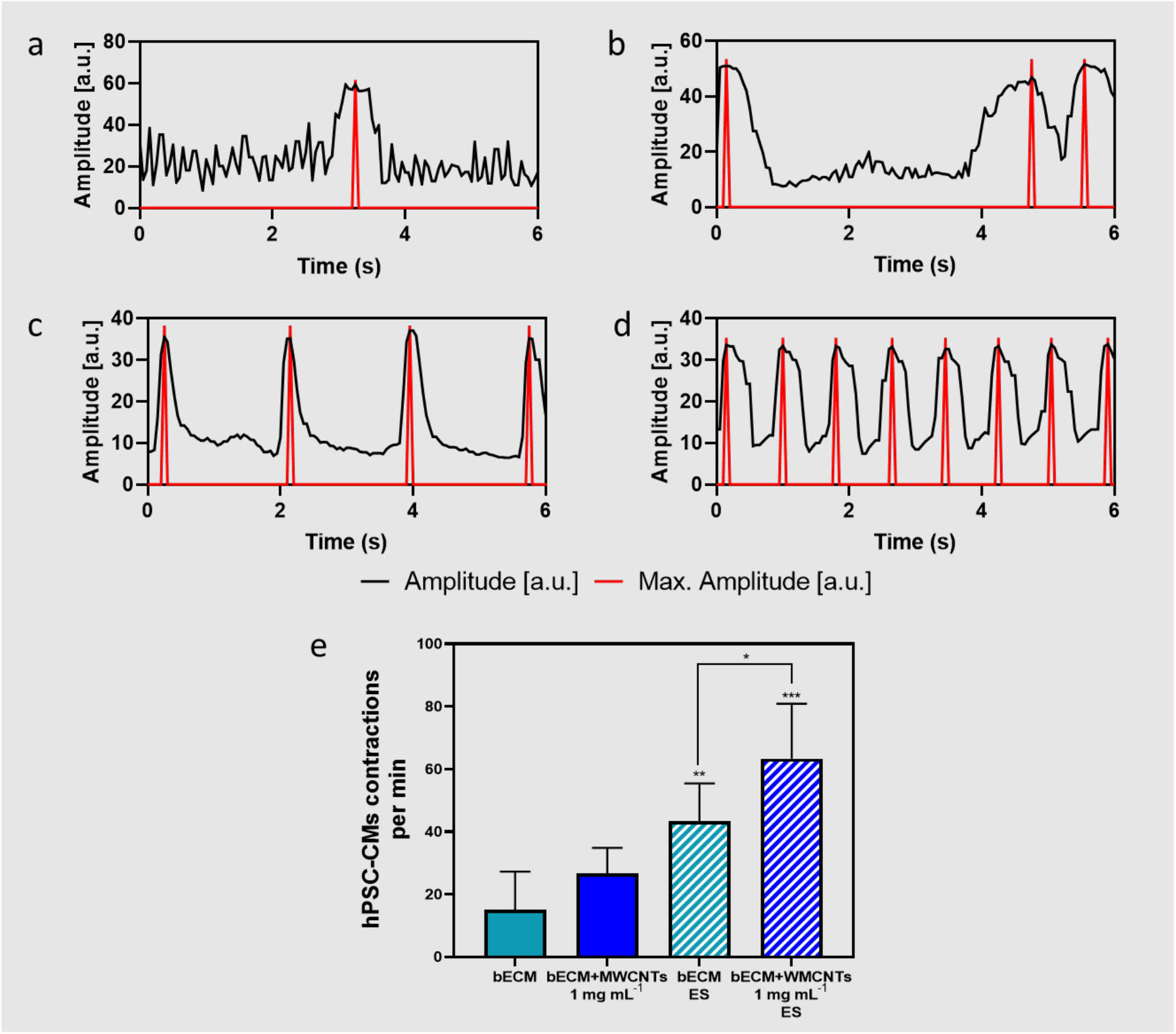
Time-dependent changes in autonomous contractile behaviour of hPSC-CMs determined using the analytical tool Myocyter (v1.3), where contractions translate to positive going transients with an arbitrary unit (a.u.), of **(a)** bECM, **(b)** bECM+MWCNTs 1mg mL^-1^, **(c)** bECM under electrical stimulation (ES) and **(d)** bECM+MWCNTs 1mg mL^-1^ under ES. **(e)** Contraction rate of hPSC-CMs per minute on the different samples (n=3) (*p = 0.0442, **p = 0.0024, ***p = 0.0009).

Once several differences in the contractile behaviour of hPSC-CMs were established, we proceeded to evaluate any effects in the genotype of the cells using markers to determine cell maturity, morphology, electrophysiology and calcium handling behaviour by RT-PCR analysis. In this case, RT-PCR was selected for this analysis as it can provide more precise information on the developmental state of cells and because fluorescence markers are difficult to visualise in cells encapsulated in 3D hydrogels.

The maturation state of hPSC-CMs was obtained from the developmentally controlled and irreversible genetic switch in the TNNI gene. The TNNI isoform switch has been frequently used as a quantitative maturation signal for hPSC-CMs [53]. The TNNI1 gene (ssTnI) is expressed in the sarcomeres of foetal and neonatal hearts, which is then replaced by the TNNI3 (cTnI) isoform. The ratio between TNNI3/TNNI1 can provide information about the maturation state of the cells. The level of TNNI3/TNNI1 expression was similar in bioprinted cells in bECM than in controls (**Figure 6a**), indicating that the levels of maturity were not significantly different. This was also the case of bioprinted hPSC-CMs in bECM+MWCNTs without ES. For bioprinted cells in bECM under ES, there is a slight increase in the TNNI3/TNNI1 expression values, however, this was not enough to be significantly different. This was not the case of hPSC-CMs bioprinted in bECM+MWCNTs under ES, where a significant difference can be observed when compared to bECM. This result suggests that the combination of the conductivity and ES of the materials can potentially increase the maturation level of the cells. Additional maturation studies based on alternative morphological and genetic markers need to be performed to confirm this hypothesis.

**Figure 6.**
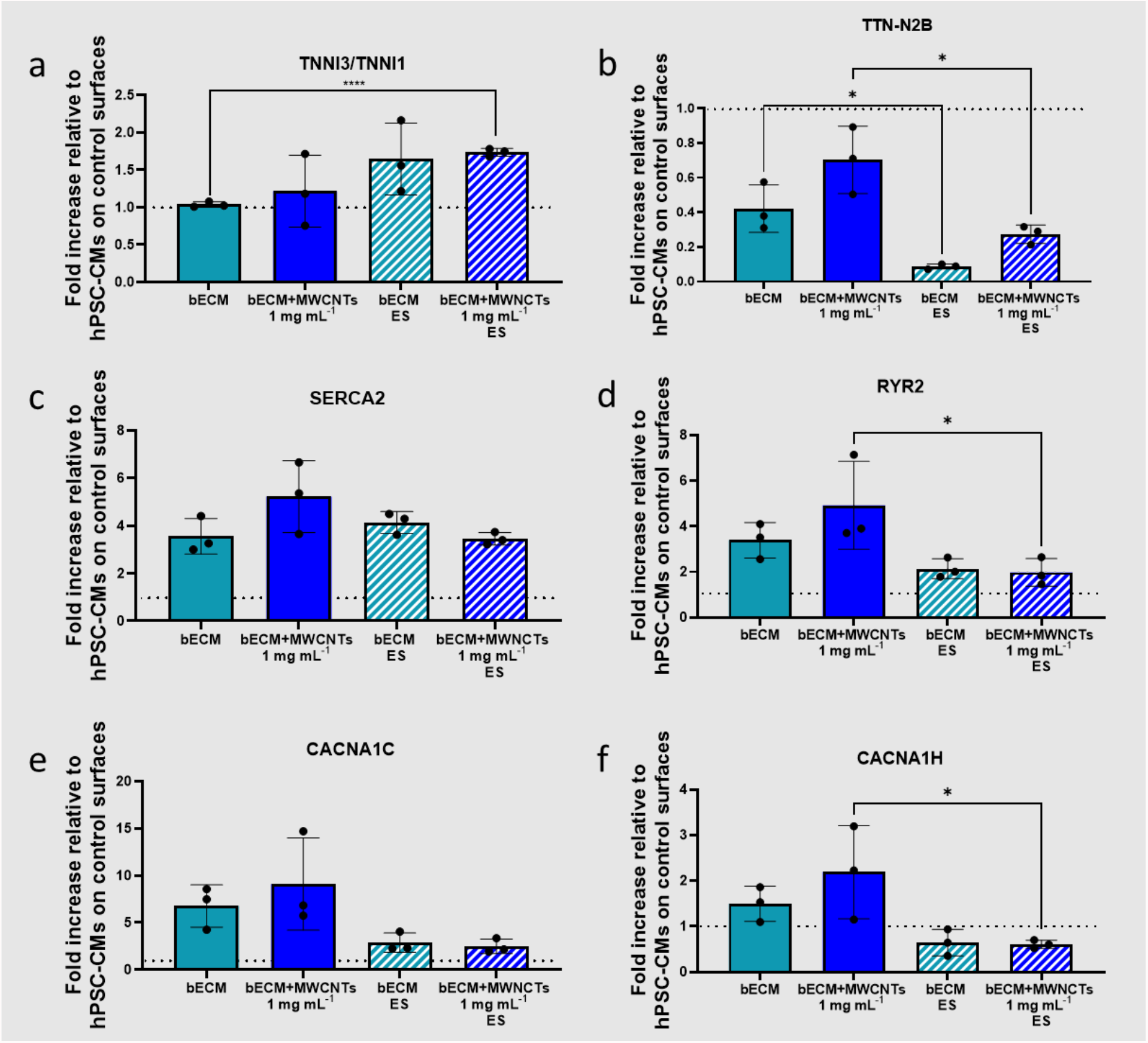
RT-qPCR analysis of the hPSC-CMs on bECM and bECM+MWCNTs 1mg mL^-1^ with and without ES. **(a)** TNNI3/TNNI1, **(b)** TTN-N2B, **(c)** SERCA2, **(d)** RYR2, **(e)** CACNA1C and **(f)** CACNA1H gene expressions are normalised against the housekeeping gene GAPDH and presented as fold-change levels relative to hPSC-CMs on control surfaces. Results expressed as mean ±SD of three (n=3) independent experiments.

TTN-N2B was selected as a marker of cell morphology and function, as this is an elastic protein expressed in the sarcomeres of hPSC-CMs. The RT-PCR results showed that this gene was downregulated in all cases when compared to controls (**Figure 6b**). We hypothesise that this could be due to the non-typical morphology of the hPSC-CMs encapsulated in the hydrogels, where a spherical morphology was observed (**Figure 2** and **Figure S7**). Future strategies will be aimed at achieving higher control over the nano- and micro-topographies of the hydrogels to enhance the morphology of the cells.

The expression of the calcium-handling genes SERCA2 and RYR2, involved in the excitation-contraction coupling, was then evaluated. Previous studies reported an increased expression of these gens in hPSC-CMs when encapsulated in 3D hydrogels of cardiac dECM [54]. In our study, these genes were upregulated in all cases (**Figure 6c-d**), however, no major differences were observed in the bioprinted structures subjected to ES. Similar results were previously reported and it has been hypothesised that although ES could induce electrophysiological alterations similar to native development, they may not promote all aspects of the complex process of electrophysiological maturation [55].

Expression of electrophysiological markers of hPSC-CMs was also evaluated through the CACNA1C and CACNA1H gene expression, coding for L-type and T-type calcium channels respectively. L-type Ca^2+^ currents contribute to the shape of the cardiac action potential, and its regulation plays an important role in cardiac excitability and contractility [56]. The CACNA1C gene was overexpressed in all cases (**Figure 6e**) and its expression was more accentuated when no ES was applied, similarly to the case of SERCA2 and RYR2. T-type Ca^2+^ currents are more present in pace making heart cells but absent in the myocardium. In the absence of ES, the CACNA1H gene was upregulated, however, when external ES was applied, this stimulation was predominant for the pacing of the cells and thus, this gene was downregulated when ES was applied (**Figure 6f**).

## 4. Conclusions

In this work, a strategy to develop novel and versatile bioinks for FRESH extrusion bioprinting is proposed with potential applications in bioelectronics. The main component of this bioink is decellularized extracellular matrix, providing a suitable environment for cell growth with tailored biochemical signalling. 3D bio-/printed structures were successfully manufactured, encapsulating hPSC-CMs while maintaining high cell viability. To further explore the sensing/actuating potential of this material, MWCNTs were dispersed in the hydrogel matrix, enhancing the electrical properties and structure of printed dECM hydrogels. Electrical stimulation was then applied and our results showed that the combination of an electrically conductive material with external electrical stimulation can drive contraction rates similar to physiological conditions. This demonstrates the potential of this material to be used in the development of smart scaffolds for biosensing/actuating applications. RT-PCR results indicated a significant improvement in the maturity markers of hPSC-CMs and downregulation of L-type Ca^2+^ currents, typically observed in pacemaker heart cells which can be an indication that the profile of the cells is more aligned to those found in the myocardium. Nevertheless, morphological features of these cells were not optimal, exhibiting a spherical shape instead of the typical rod shape of the cardiomyocytes probably due to the lack of resolution of the extrusion bioprinting technique. Further development in the 3D bioprinting technology, combining techniques that can accurately reproduce the macro-, micro- and nano- architectures of tissues are required to achieve higher biomimicry.

## Acknowledgements

We would like to thank Professor Michael McAlpine for support, guidance and providing access to laboratory facilities and training of Dr Paola Sanjuan-Alberte whist visiting the University of Minnesota. The University of Nottingham is acknowledged for supporting the EPSRC Doctoral Prize award granted to PSA. Part of this work was carried out in the Characterization Facility, University of Minnesota, which receives partial support from NSF through the MRSEC program.

## Author Contributions

PSA contributed to the tissue decellularization and bioink formulation established the extrusion bioprinting protocols and characterisation of the printed structures and performed hPSC-CMs differentiation and assays. CW performed the sisECM decellularization and characterisation and rheological studies. JNJ and SK performed bECM and lECM decellularization and characterisation. JCS contributed towards the primers’ selection, RNA extraction and RT-PCR analysis. NC contributed to the sample preparation and analysis by SEM. PSA, LJW and FJR contributed to the project conceptualisation. PSA, RJMH, JMSC, FCF, LJW and FJR contributed to the experimental design, analysis and interpretation of results. All authors read and approved the final manuscript.

