## Supporting information for "Bio-inspired artificial printed bioelectronic cardio-3D-cellular constructs"

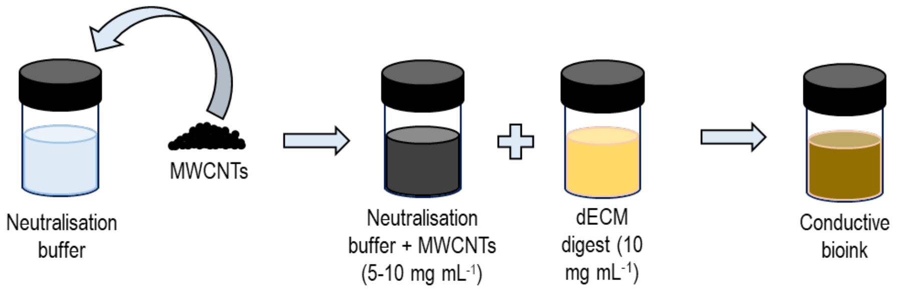


**Figure S1.** Schematic showing the procedure for preparation of the conductive bioinks

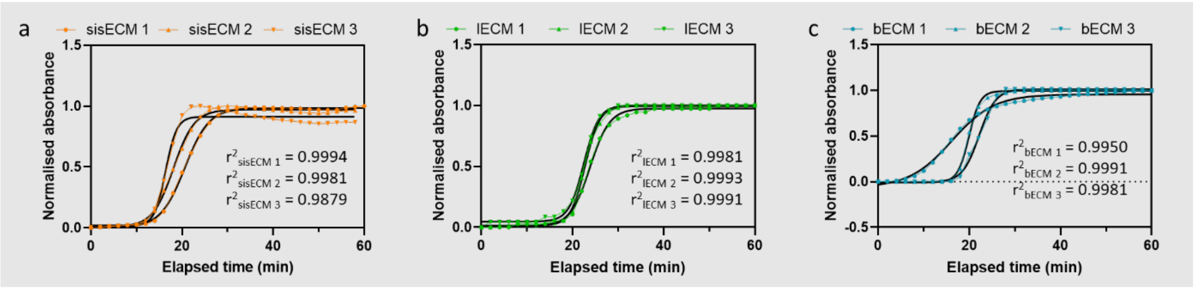


**Figure S2**. Fitting of sigmoidal curves from the normalised absorbance values obtained during the gel kinetics determination of (a) sisECM, (b) lECM and (c) bECM at 450 nm.

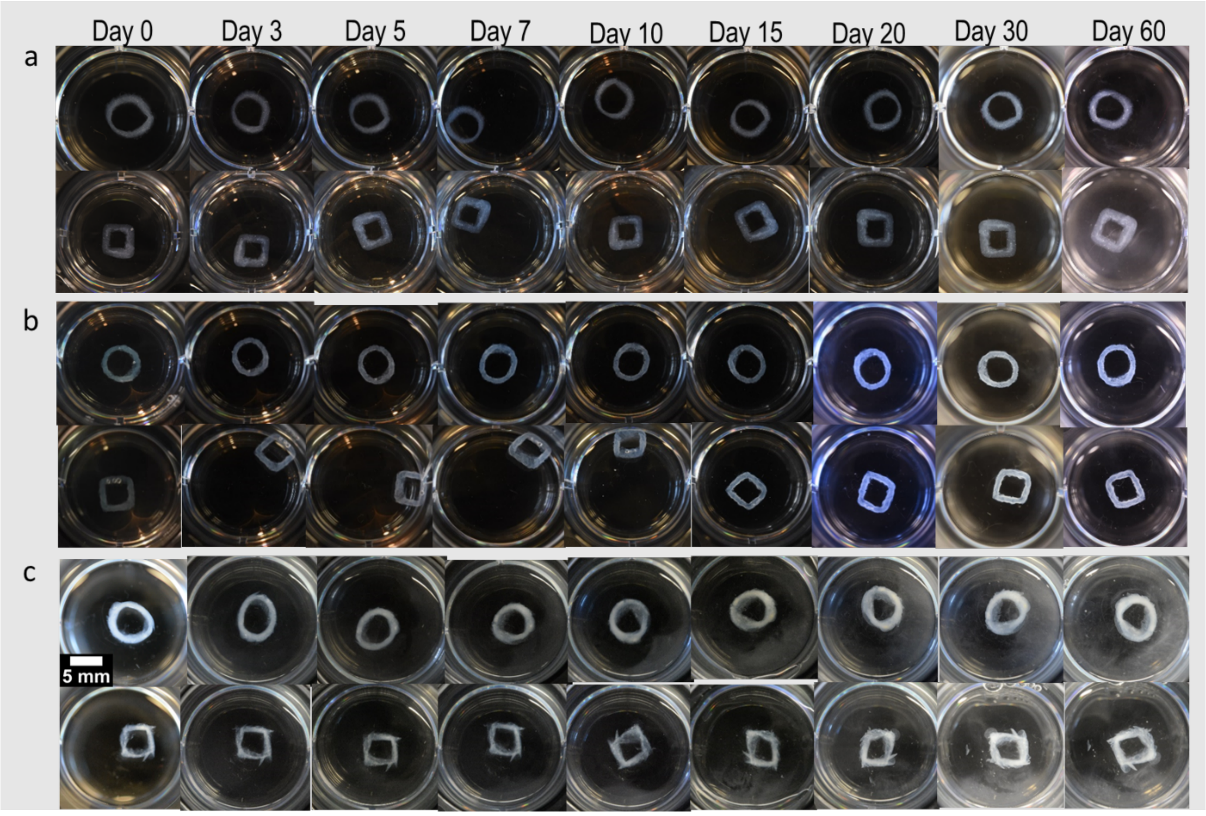


**Figure S3.** Stability of FRESH extruded **(a)** sisECM, **(b)** lECM and **(c)** bECM structures over time. These structures consisted on 6 mm diameter rings and squares with 6 mm sides.

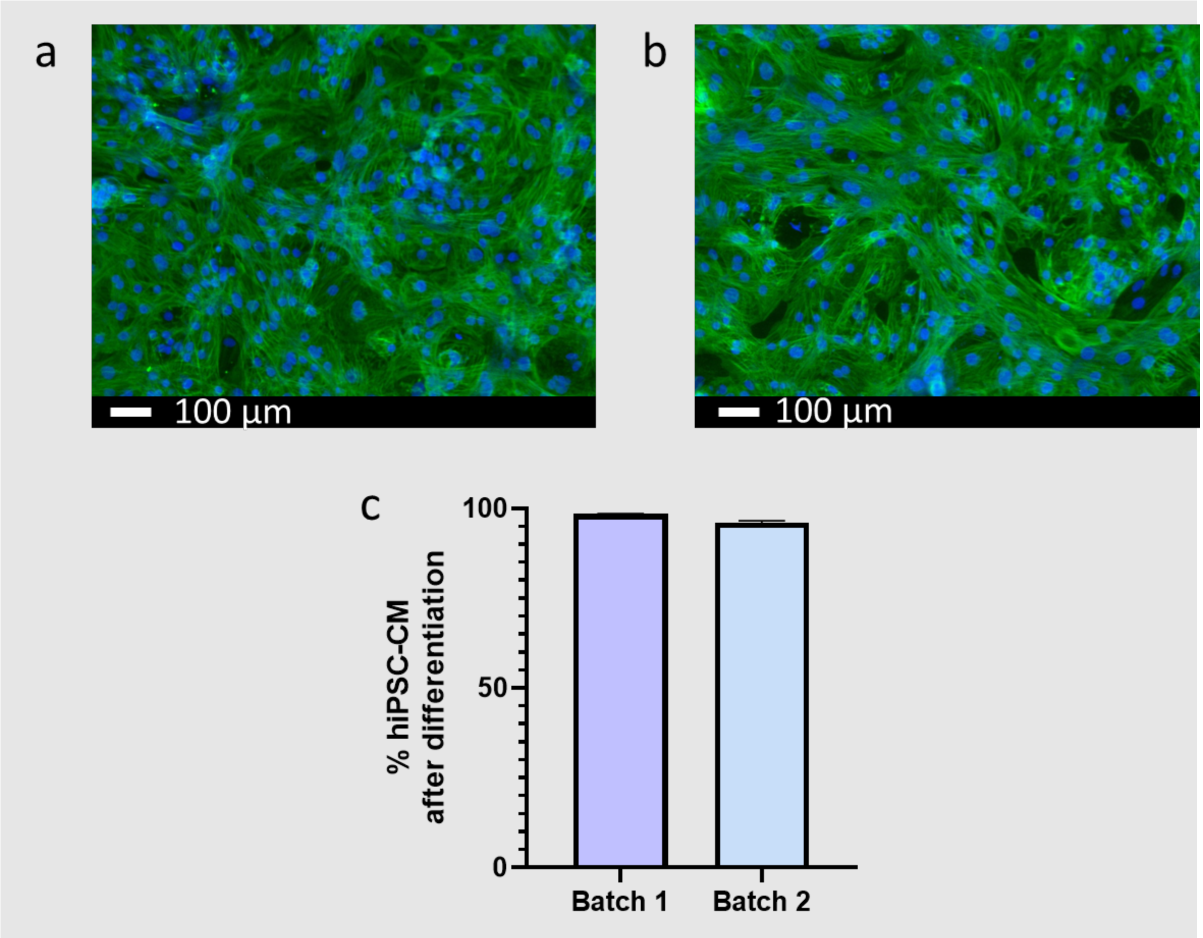


**Figure S4.** Immunostaining of two different batches **(a,b)** of hPSC-CMs after differentiation. Representative images were selected. Cells were immunolabelled for cardiac troponin TNNI3 (green). Nuclei were counterstained with Hoechst (blue). hPSC-CMs are positive for the green markers, whereas other cell types are stained only with Hoechst. **(c)** Determination of hPSC-CMs purity of the differentiated batches.

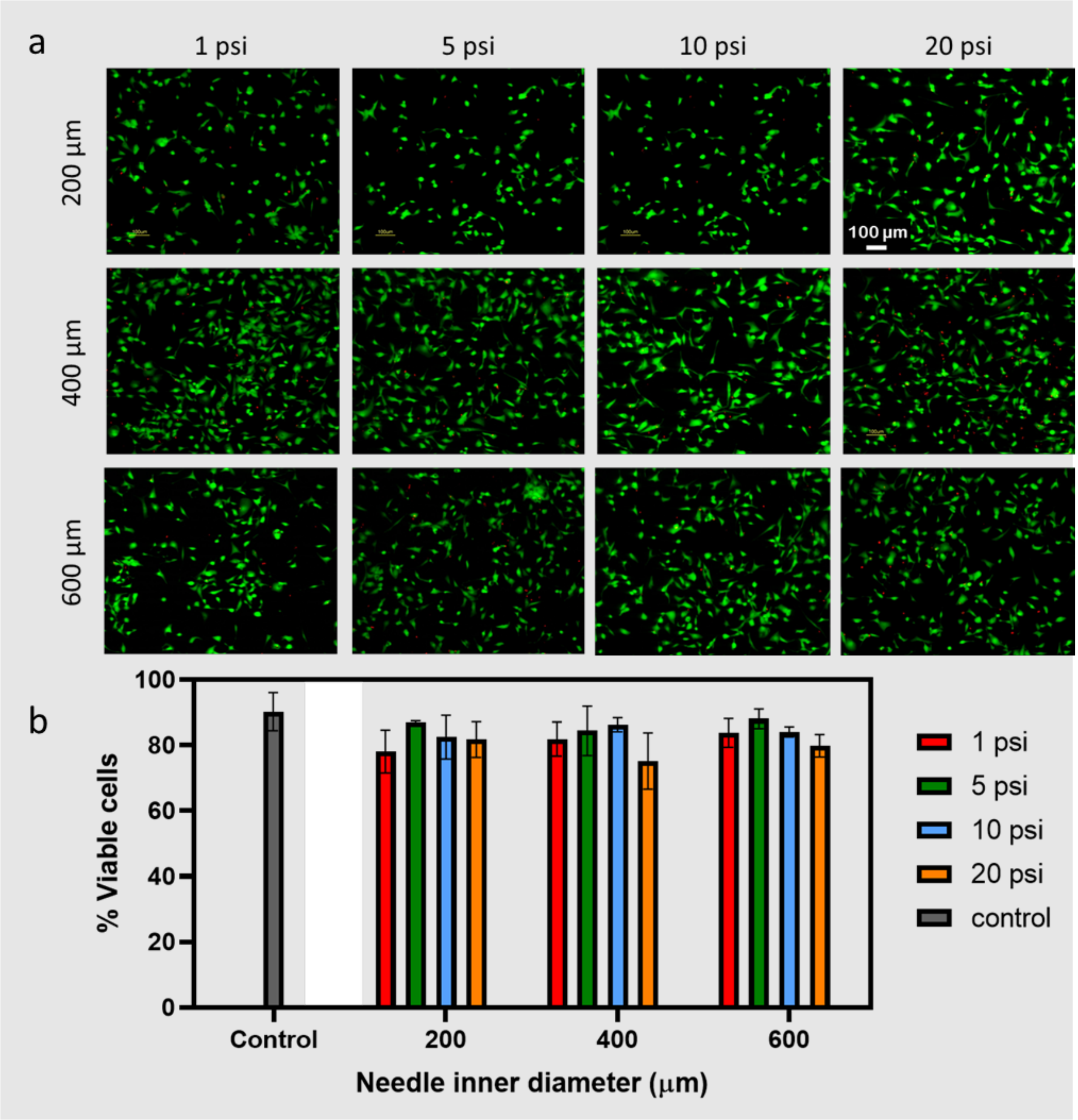


**Figure S5. (a)** Live/Dead staining of human pluripotent stem cell derived cardiomyocytes (hPSC-CMs) 24 hours after bioprinting using various inner needle diameters and pressures. Scale bar 100 µm. **(b)** Calculated percentage of viable cells for each condition (n=3, ±SD).

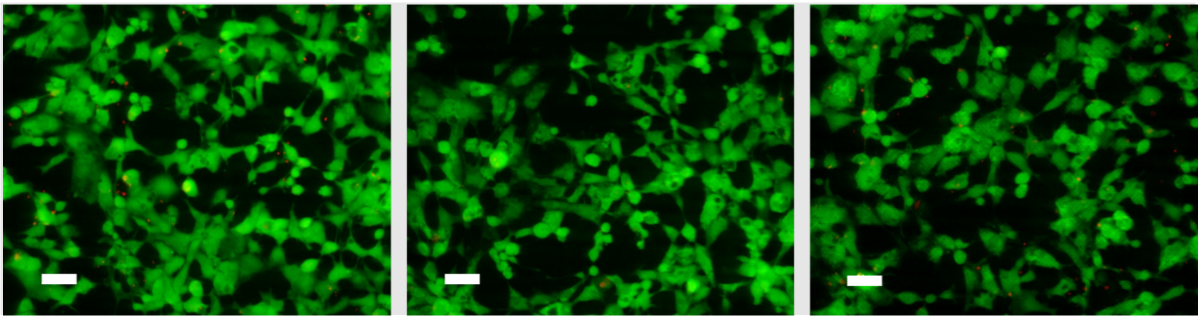

**Figure S6**. Live/Dead staining of hPSC-CMs in three different samples cultured on control surfaces (Well-plate). Scale bar 50 µm

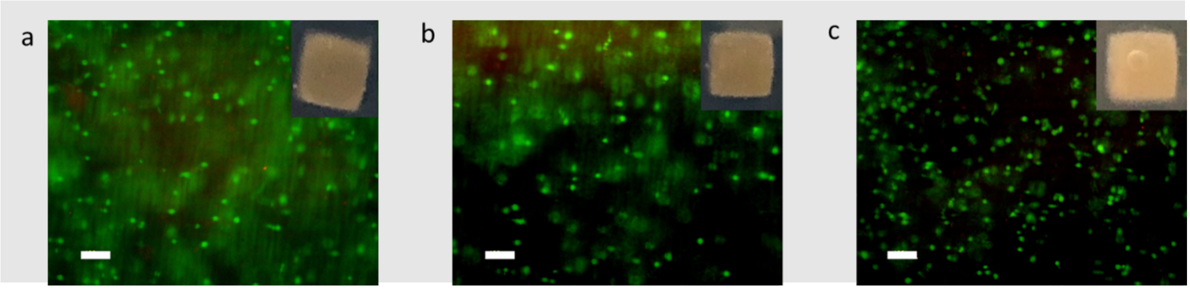


**Figure S7**. Live/Dead staining of hPSC-CMs encapsulated on **(a)** sisECM+MWCNTs 1 mg ml$^{-1}$, **(b)** lECM+MWCNTs 1 mg ml$^{-1}$, **(c)** bECM+MWCNTs 1 mg ml$^{-1}$ and inset images of the different hydrogels. Scale bar 100 μm.

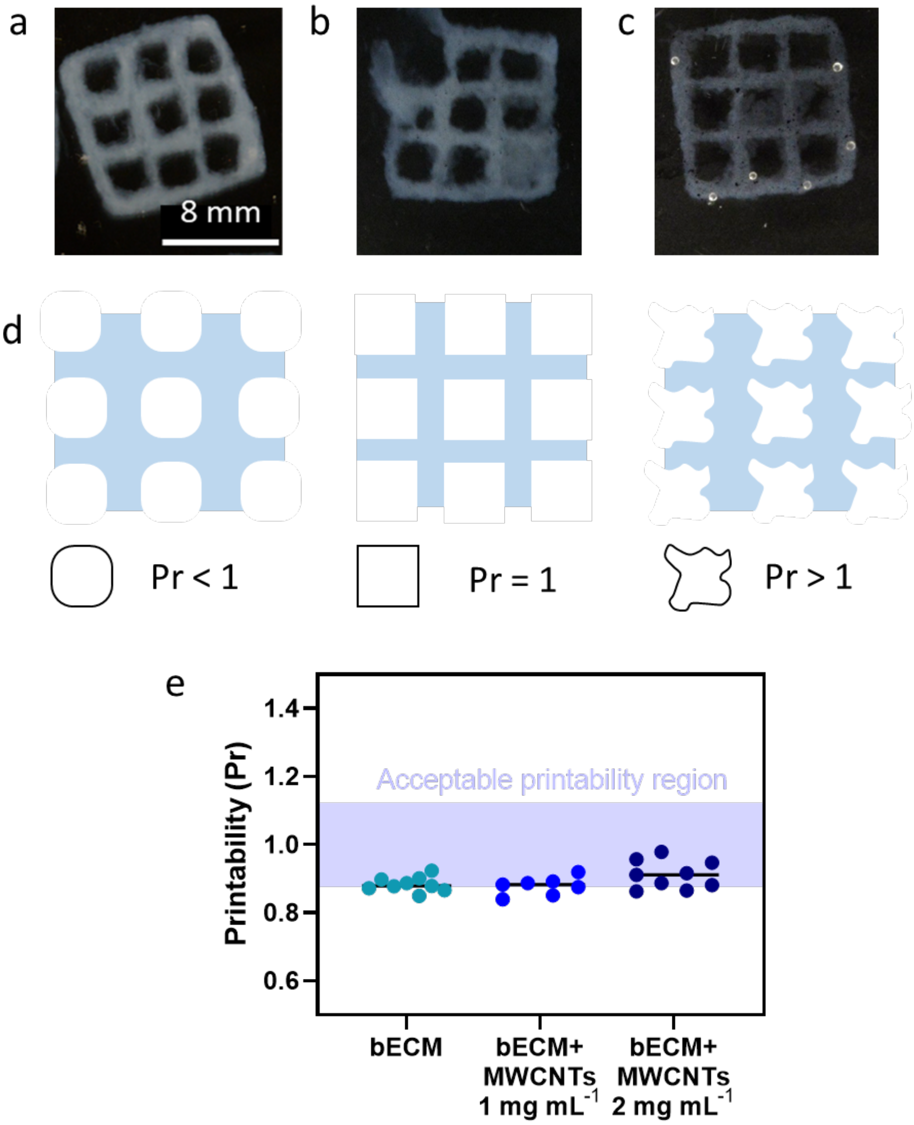


**Figure S8.** Printed 10 mm square scaffold using (a) bECM, (b) bECM+MWCNTs 1 mg mL-1 and (c) bECM+MWCNTs 2 mg mL-1. (d) Evaluation of printability (Pr) under three typical conditions. (e) Semi-quantified Pr value of printed constructs.

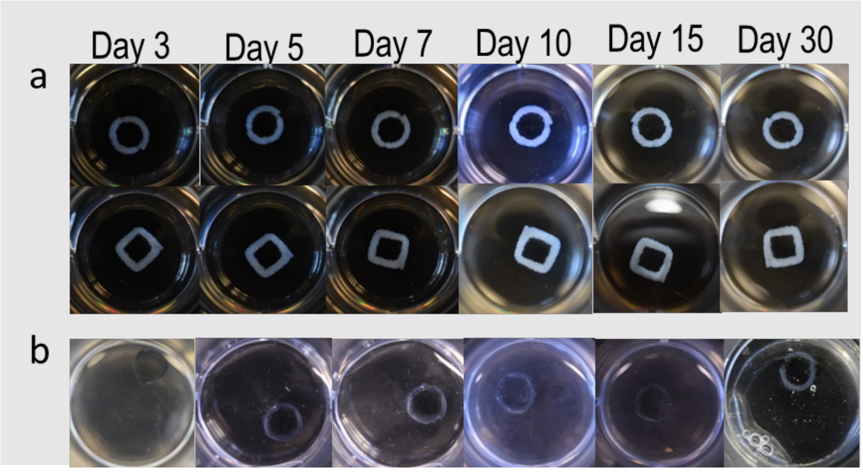


**Figure S9.** Stability of FRESH extruded bECM at **(a)** 1 mg $mL^{-1}$ and **(b)** 2 mg $mL^{-1}$ multi-walled carbon nanotubes (MWCNTs) concentration.

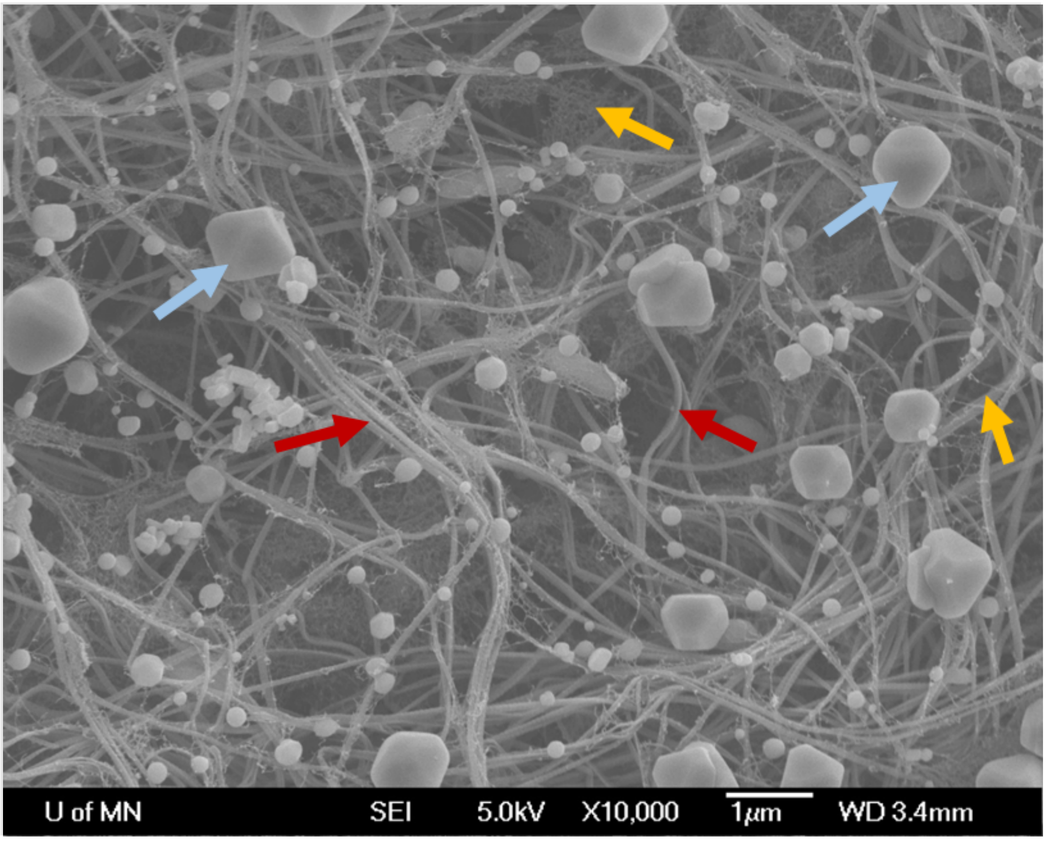


**Figure S10.** SEM images of bECM and MWCNTs at a concentration of 0.2 mg $mL^{-1}$ the different elements can be observed: collagen fibres (red arrows), MWCNTs (yellow arrows) and buffer salts (blue arrows).

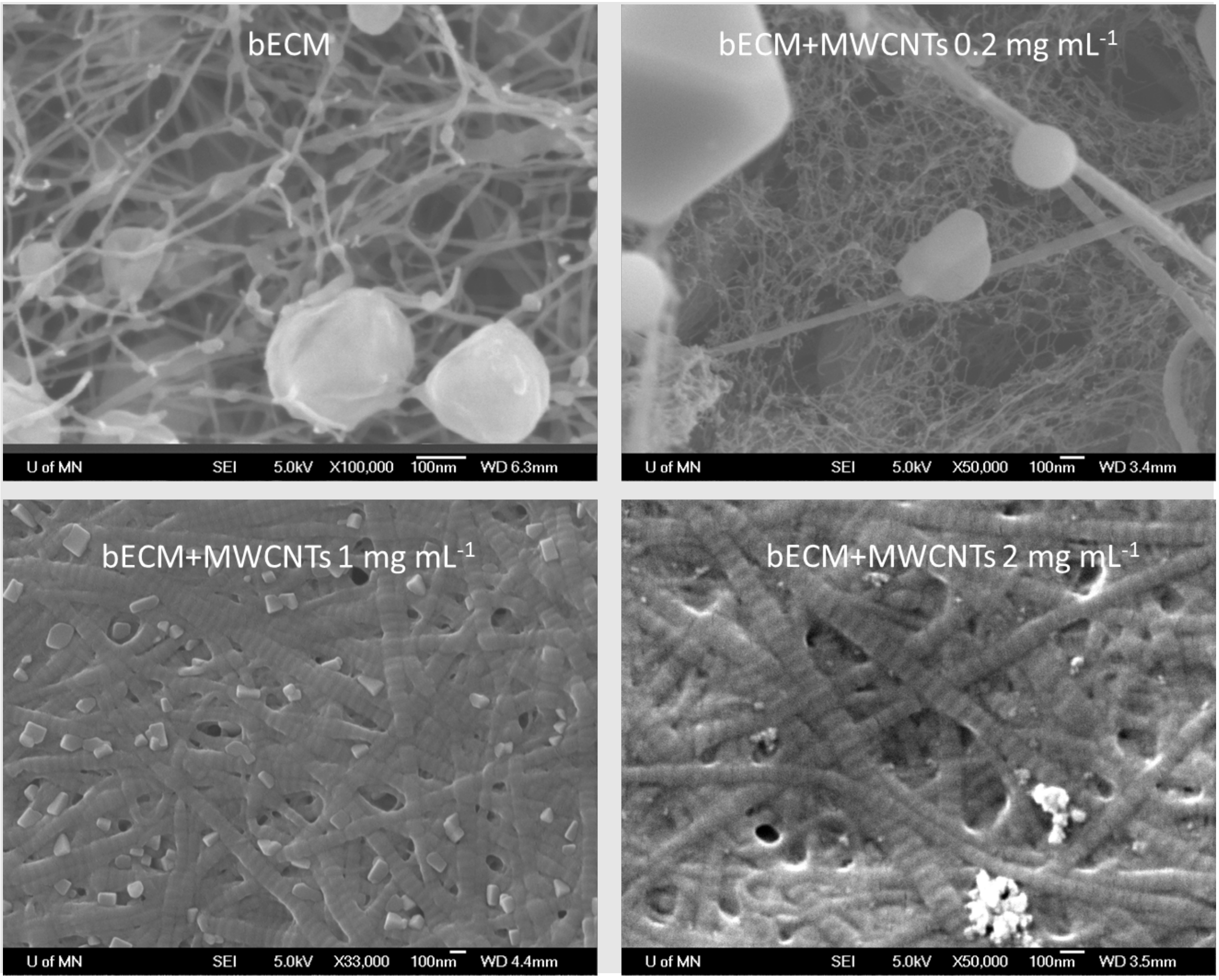


**Figure S11**. SEM images at higher magnifications

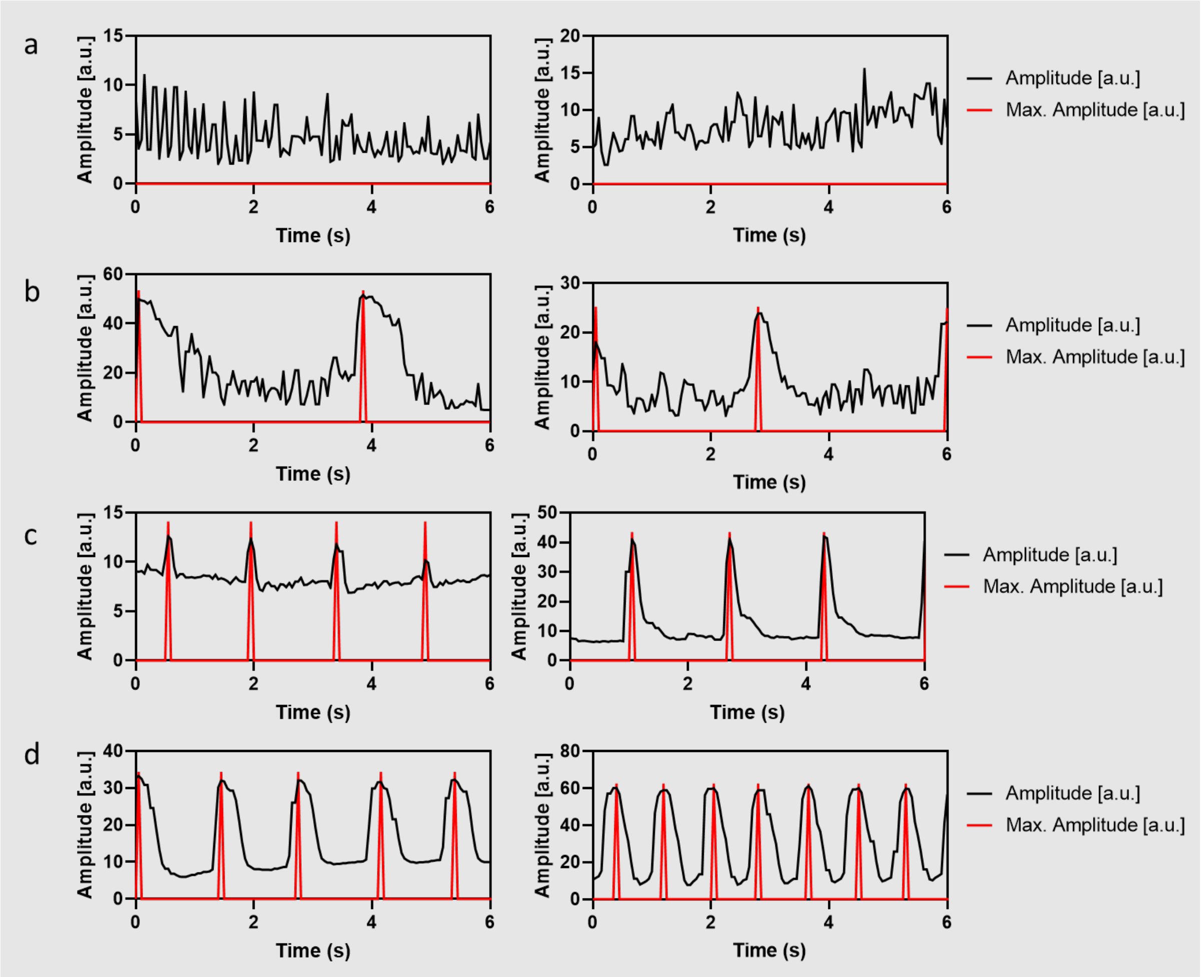


**Figure S12.** Time-dependent changes in autonomous contractile behaviour of hPSC-CMs of additional replicates determined using the analytical tool Myocyter (v1.3) of (a) bECM, (b) bECM+MWCNTs 1mg mL-1, (c) bECM under electrical stimulation (ES) and (d) bECM+MWCNTs 1mg mL-1 under ES.

**Table S1**. List of primers used

| Gene | | Sequence | Size (bp) |
|---|---|---|---|
| cTNNI (TNNI3) | Forward | CCTCCAACTACCGCGCTTAT | 20 |
| | Reverse | CTGCAATTTTCTCGAGGCGG | 20 |
| ssTNNI (TNNI1) | Forward | GCTCCACGAGGACTGAACAA | 20 |
| | Reverse | CTTCAGCAAGAGTTTGCGGG | 20 |
| TTN-2NB | Forward | CCAATGAGTATGGCAGTGTCA | 21 |
| | Reverse | TACGTTCCGGAAGTAATTTGC | 21 |
| SERCA2 | Forward | ACCCACATTCGAGTTGGAAG | 20 |
| | Reverse | CCAACGAAGGTCAGATTGGT | 20 |
| RYR2 | Forward | AAGCCCTCTCGTCTGAAACA | 20 |
| | Reverse | CCACCCAGACATTAGCAGGT | 20 |
| CACNA1C | Forward | CAATCTCCGAAGAGGGGTTT | 20 |
| | Reverse | TCGCTTCAGACATTCCAGGT | 20 |
| CACNA1H | Forward | TCAACGTCATCACCATGTCC | 20 |
| | Reverse | AGCCTCGAAGACAAACACGA | 20 |
| GAPDH | Forward | AACAGCGACACCCACTCCTC | 20 |

| | | | |
|---|---|---|---|
| | Reverse | CATACCAGGAAATGAGCTTGACAA | 24 |

**Video S1.** hPSC-CMs contraction rate on bECM and no electrical stimulation

**Video S2.** hPSC-CMs contraction rate on bECM+MWCNTs 1 mg $mL^{-1}$ and no electrical stimulation

**Video S3.** hPSC-CMs contraction rate on bECM after electrical stimulation

**Video S4.** hPSC-CMs contraction rate on bECM+MWCNTs 1 mg $mL^{-1}$ after electrical stimulation
